# Auditory Motion Does Not Modulate Spiking Activity in the Middle Temporal and Medial Superior Temporal Visual Areas

**DOI:** 10.1101/204529

**Authors:** Tristan A. Chaplin, Benjamin J. Allitt, Maureen A. Hagan, Marcello G.P. Rosa, Ramesh Rajan, Leo L. Lui

**Author notes:** Correspondence should be addressed to: Tristan Chaplin, 26 Innovation Walk, Monash University, Victoria, Australia, 3800, and Leo Lui, 26 Innovation Walk, Monash University, Victoria, Australia, 3800,.

## Abstract

The integration of multiple sensory modalities is a key aspect of brain function, allowing animals to take advantage of concurrent sources of information to make more accurate perceptual judgments. For many years, multisensory integration in the cerebral cortex was deemed to occur only in high-level “polysensory” association areas. However, more recent studies have suggested that cross-modal stimulation can also influence neural activity in areas traditionally considered to be unimodal. In particular, several human neuroimaging studies have reported that extrastriate areas involved in visual motion perception are also activated by auditory motion, and may integrate audio-visual motion cues. However, the exact nature and extent of the effects of auditory motion on the visual cortex have not been studied at the single neuron level. We recorded the spiking activity of neurons in the middle temporal (MT) and medial superior temporal (MST) areas of anesthetized marmoset monkeys upon presentation of unimodal stimuli (moving auditory or visual patterns), as well as bimodal stimuli (concurrent audio-visual motion). Despite robust, direction selective responses to visual motion, none of the sampled neurons responded to auditory motion stimuli. Moreover, concurrent moving auditory stimuli had no significant effect on the ability of single MT and MST neurons, or populations of simultaneously recorded neurons, to discriminate the direction of motion of visual stimuli (moving random dot patterns with varying levels of motion noise). Our findings do not support the hypothesis that direct interactions between MT, MST and areas low in the hierarchy of auditory areas underlie audiovisual motion integration.

## Introduction

The natural environment often produces stimuli that can be perceived by multiple senses, making multisensory integration one of the fundamental aspects of brain function (Stein & Stanford, 2008; Stein *et al.*, 2014). There is evidence that humans and monkeys can integrate multisensory cues in a statistically optimal way (Ernst & Banks, 2002; Gu *et al.*, 2008; Parise *et al.*, 2012), giving a more reliable account than either sense alone.

Many studies of the neurological basis of multisensory integration have been shaped by a model of sensory processing in which each sensory modality is processed independently and only integrated in higher-level cortical areas (Felleman & Van Essen, 1991; Wallace *et al.*, 2004) or in subcortical structures that receive converging projections from unisensory areas of different modalities (Meredith and Stein 1983; Meredith et al. 1987; Reig and Silberberg 2014). However, studies have now shown that low level sensory cortical areas, historically considered unisensory, can be influenced by other modalities (for a review see Ghazanfar and Schroeder, 2006). In particular, these studies have shown cross-modal influences in the auditory and visual systems (Schroeder & Foxe, 2002; Bizley *et al.*, 2007; Lakatos *et al.*, 2007; Bizley & King, 2008, 2009; Wang *et al.*, 2008; Iurilli *et al.*, 2012; Olcese *et al.*, 2013; Meijer *et al.*, 2017).

Studies of visual motion perception have provided some of the most in-depth insights into the relationship between sensory neurons and behavior (Parker and Newsome, 1999), and motion stimuli have also proven to be a useful tool for understanding multisensory integration in low level visual areas (Gu et al., 2008; Fetsch et al., 2013). Although it is well know that visual motion processing areas can integrate self-motion cues from the vestibular system (Duffy, 1998; Gu *et al.*, 2006), the integration of moving audio-visual cues has remained more controversial. Some human imaging studies have reported that the addition of a moving sound to a moving visual stimulus can modulate the responses in the human visual motion processing complex (hMT+, Alink et al., 2008; Lewis and Noppeney, 2010; von Saldern and Noppeney, 2013). Two imaging studies have also found evidence of responses to auditory motion alone in hMT+ (Poirier et al., 2005; Strnad et al., 2013 but see Jiang et al., 2014). The integration of audio-visual cues in hMT+ is an attractive proposition, given that psychophysical studies have shown that moving auditory cues can be integrated with visual stimuli to improve visual motion detection (Meyer & Wuerger, 2001; Kim *et al.*, 2012), that auditory motion can improve learning in visual motion tasks (Seitz *et al.*, 2006), and that auditory stimuli can influence visual motion perception (Sekuler *et al.*, 1997; Meyer & Wuerger, 2001; Kitagawa & Ichihara, 2002; Beer & Roder, 2004; Soto-Faraco *et al.*, 2005; Freeman & Driver, 2008; Alink, Euler, Galeano, *et al.*, 2012; Kafaligonul & Stoner, 2012; Kafaligonul & Oluk, 2015).

In contrast, other studies suggest that integration of audio-visual motion cues may only occur in higher-level brain areas. Some psychophysical studies (Wuerger *etal*, 2003; Alais & Burr, 2004) have found that observer performance is consistent with probability summation, i.e. the improved performance results from the fact that observers have two chances (i.e. visual or auditory) to answer correctly. This type of integration is likely to occur in a high-level brain region (Bizley *et al.*, 2016), and human imaging studies often report activity in such regions (Lewis *et al.*, 2000; Baumann & Greenlee, 2007; von Saldern & Noppeney, 2013).

To test if neurons in visual motion processing areas of the primate cerebral cortex integrate auditory motion cues without the influence of top-down pathways, we performed extracellular recordings in the middle temporal (MT) and medial superior temporal (MST) areas (Figure 1A), which together comprise the homolog of hMT+ (Zeki *et al.*, 1991; Huk *et al.*, 2002), in anaesthetized marmosets. In marmosets, these areas receive sparse connections from auditory cortex (Palmer & Rosa, 2006a, 2006b), and their human homologs have been implicated in the perception of moving auditory stimuli (Baumgart & Gaschler-Markefski, 1999; Pavani *et al.*, 2002; Warren *et al.*, 2002; Ducommun *et al.*, 2004; Alink, Euler, Galeano, *et al.*, 2012). We tested two hypotheses: first, that neurons in MT and MST show responses to unimodal auditory stimuli, and second, that concurrent auditory and visual stimulation in the same direction of motion facilitates neuronal responses, in comparison with visual motion only. The second hypothesis was tested with levels of visual noise, as stimulus conditions that are more difficult to discriminate often reveal more robust multisensory integration effects (Meredith & Stein, 1983; Deneve *et al.*, 2001; Ma *et al.*, 2006; Gu *et al.*, 2008; Fetsch *et al.*, 2011).

**Figure 1:**
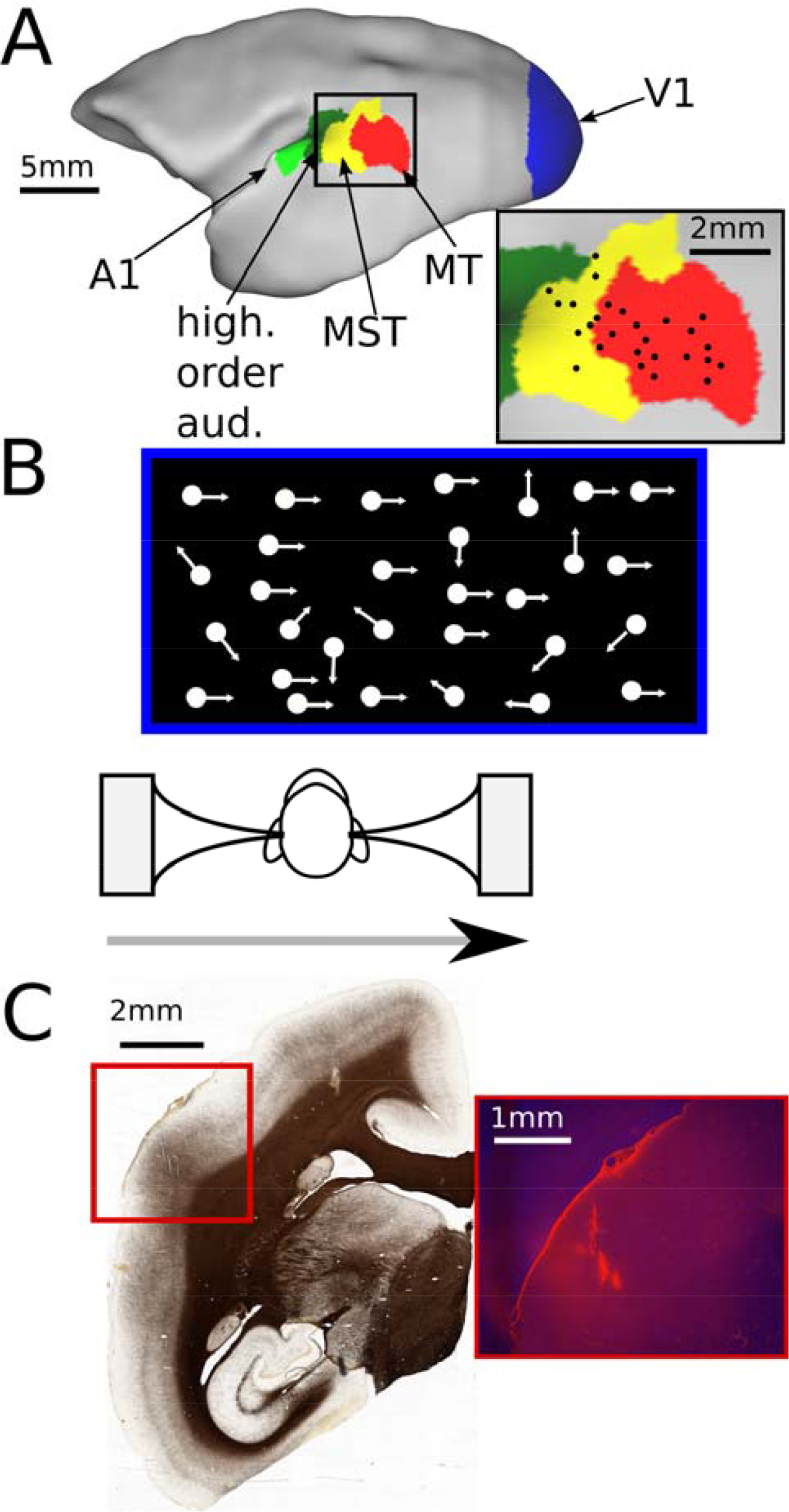
Experimental design and setup. A: Lateral view of a marmoset cerebral cortex model, with the relevant cortical areas - MT, MST, caudal higher-order auditory cortex, primary auditory cortex (A1) and primary visual cortex V1, shown for reference). The inset shows a summary map of the approximate locations of all penetrations, showing good coverage of both MT and MST. B: Experimental setup. A computer screen was positioned in front of the animal was used to present visual dot motion, either left or right, and congruent auditory motion was simulated through headphones, shown by the grey arrow. C: Histology. The image on the left shows a coronal myelin section from one case. Area MT is identifiable by the densely myelinated region in the red box. The image on the right shows the region in the red box as a fluorescence image, showing the fluorescent tracks made by the DiI-coated electrodes in area MT.

## Methods

### Animals and surgical preparation

Single-unit and multi-unit extracellular recordings in areas MT and MST were obtained from 5 marmoset monkeys (2 male and 3 female, between 1.5 and 3 years of age, with no history of veterinary complications). These animals were also used for unrelated anatomical tracing and visual physiology experiments. Experiments were conducted in accordance with the Australian Code of Practice for the Care and Use of Animals for Scientific Purposes, and the Principles and Guidelines for the Care and Use of Non-Human Primates for Scientific Purposes (National Health and Medical Research Council, 2016, Part A, pages 4-9, www.nhmrc.gov.au/guidelines/publications/ea15978-1-925129-68-7), and all procedures were approved by the Monash University Animal Ethics Experimentation Committee, which also monitored the health and wellbeing of the animals throughout the experiments. The Principles and Guidelines also governed the way which animals were housed and cared for prior to experiments (National Health and Medical Research Council, 2016, section B.5 pages 10-12); animals were group housed in large cages and received regular outdoor access, as well as receiving daily care from staff.

Anesthesia was induced with alfaxalone (Alfaxan, 8 mg/kg), allowing a tracheotomy, vein cannulation and craniotomy to be performed. After all surgical procedures were completed, the animal was administered an intravenous infusion of pancuronium bromide (0.1 mg/kg/h) combined with sufentanil (6-8 μg/kg/h, adjusted to ensure no physiological responses to noxious stimuli) and dexamethasone (0.4 mg/kg/h), and was artificially ventilated with a gaseous mixture of nitrous oxide and oxygen (7:3). The electrocardiogram and level of cortical spontaneous activity were continuously monitored. Administration of atropine (1%) and phenylephrine hydrochloride (10%) eye drops was used to produce mydriasis and cycloplegia. Appropriate focus and protection of the corneas from desiccation were achieved by means of hard contact lenses selected by retinoscopy. This preparation has been used many times to record spiking activity in the visual cortex including area MT (e.g. Lui et al., 2007) and robust spiking responses in the auditory cortex (Rajan *et al.*, 2013) including sustained responses and both “on” and “off” components of vocalizations, similar to those found in awake preparations, and sensitivity to interaural level differences (Lui et al. 2015).

### Electrophysiology, data acquisition and pre-processing

We recorded neural activity with single shaft linear arrays (NeuroNexus) consisting of 32 electrodes separated by 50 μm. MT and MST recording sites were identified during experiments using anatomical landmarks, receptive field progression and size (Rosa & Elston, 1998), and direction selectivity, and were confirmed postmortem by histological examination.

Electrophysiological data were recorded using a Cereplex system (Blackrock Microsystems) with a sampling rate of 30 kHz. For online analysis of spiking activity, each channel was high-pass filtered at 750 Hz and spikes were initially identified based on threshold crossings (-4 standard deviations of the root mean square). Threshold crossings were sorted for offline analysis using Offline Sorter (Plexon Inc.). Threshold crossings were classified as single-units if they showed good separation on the (2 component) principal component analysis plot, and were confirmed by inspection of the inter-spike interval histogram and consistency of waveform over time. Any remaining threshold crossings were classified as multi-unit activity. We excluded five single-units from adjacent channels since it was apparent they were duplicated across two channels, based on their sharp cross correlogram peak and high signal correlation (Bair *et al.*, 2001). The median spontaneous firing rate (to a blank, black screen) of multi-units was 1.9 spikes/s, indicating that the multi-units consisted of relatively few single-units. For analysis of evoked potentials, each channel was low-pass filtered at 250Hz and then filtered with a 50Hz notch filter to remove line noise.

### Visual stimuli

Visual stimuli were presented on a VIEWPixx3D monitor (1920 × 1080 pixels; 520 × 295 mm; 120 Hz refresh rate, VPixx Technologies) positioned 0.35 to 0.45 m from the animal on an angle to accommodate the size and eccentricity of the receptive field(s), typically subtending 70° in azimuth, and 40° in elevation. All stimuli were generated with MATLAB using Psychtoolbox-3 (Brainard, 1997).

The main visual stimulus consisted of random dots presented full screen. White dots (106 cd/m^2^) of 0.2° in diameter were displayed on a black (0.25 cd/m^2^) background (full contrast). The density was such that there were on average 0.5 dots per °^2^; this was chosen because these parameters elicit good responses from marmoset MT when displayed on LCD monitors (Solomon *et al.*, 2011; Zavitz *et al.*, 2016). Dot coherence was controlled using the white noise method (i.e. Britten et al., 1992, 1996; see Pilly and Seitz 2009) by randomly choosing a subset of “noise” dots on each frame, which were displaced to random positions within the stimulus aperture. The remaining “signal” dots were moved in the same direction with a fixed displacement. We also presented an 8ms full screen flash stimulus at 1Hz (100 repeats) to test for visually evoked potentials. Robust direction selective spiking responses have been demonstrated for MT neurons in responses to these stimuli in the same preparation (Chaplin *et al.*, 2017).

### Auditory stimuli

The ear canals were surgically exposed to allow the insertion of sound delivery tubes connected to speakers (MF1, Tucker-Davis Technologies). This method has the advantage of bypassing the outer ear, where certain frequencies are attenuated, ensuring that auditory stimuli will be delivered to the inner ear with greater precision and reliability. It has been used previously for evoking auditory spiking responses in the cortex of anaesthetized marmosets (Rajan *et al.*, 2013; Lui *et al.*, 2015) as well as cats (Rajan, 2000). The sound pressure level (SPL) and frequency response function of each speaker was calibrated using a Type 2673 microphone (Bruel & Kjaer) connected to a sound level calibrator type 4230 (94 dB-1000 Hz, Bruel & Kjaer), powered by a type 2804 microphone power supply. Experiments were conducted in a sound attenuating room.

The main auditory stimulus was a 6-12kHz band pass noise stimulus, chosen to match the common marmoset vocalization range (Agamaite *et al.*, 2015), which was created with an 11th order Butterworth filter, presented at 70dB average binaural intensity (ABI). At this intensity and frequency range, auditory stimuli can be easily detected by humans and marmosets (Osmanski & Wang, 2011). Similar intensity levels have been used to investigate audiovisual integration in humans (Meyer & Wuerger, 2001; Kim *et al.*, 2012) and induced BOLD responses hMT+ in fMRI studies (von Saldern & Noppeney, 2013). This stimulus was randomly generated for each trial. We used this type of stimulus as both a stationary (200 ms) and moving (500-1000 ms) sound to test for spiking responses to auditory stimuli. We varied the apparent spatial position of the static stimuli, and the direction of the moving stimuli, by manipulating the interaural level difference (ILD), the main cue for high frequency sounds in the azimuth in marmosets (Slee & Young, 2010). This approach allowed us to present moving auditory and visual stimuli concurrently, as a moving speaker would either block the visual stimulus on the monitor, or the monitor would cast an acoustic shadow on the speaker.

The stationary stimuli were presented at ILDs ranging from −25 dB to +25 dB in 5 dB increments. We used published data on the head related transfer function (Slee & Young, 2010) to guide the modulation of the ILD for the moving stimulus. For this frequency range, the ILD is approximately 5 dB at 15° azimuth, 10 dB at 30° azimuth and 15 dB at 90° azimuth. The moving auditory stimulus moved at a speed of 60°/s from the midline (0 dB ILD) to 36° azimuth (~10dB ILD, Slee & Young 2010) and vice versa for the opposite direction of motion. Cells in the marmoset auditory cortex under the same preparation are sensitive to ILD differences as little as 5 dB for pure tones and broadband stimuli at a range of intensity levels (30-70 dB ABI), including levels used in the current experiment (Lui *et al.*, 2015). We also presented auditory clicks (0.1 ms square wave, 70dB SPL) at 1Hz to test for auditory evoked potentials (100 repeats).

### Audio-visual motion stimuli

Given that the auditory motion stimuli were produced by modulating the ILD, we could only simulate auditory motion in the horizontal plane. Thus we presented visual and auditory motion moving leftwards or rightwards (Figure 1B). For visual stimuli, we presented dots at coherences of 100, 82, 64, 46, 28, 10 and 0%. For auditory stimuli, there were no such coherence conditions, only leftwards or rightwards motion. In audiovisual conditions, the direction of auditory motion was the same as the visual motion, except in the case of 0% coherence visual motion, which we presented with both left and right auditory motion. Both the visual and auditory stimuli were presented for 600ms, with 60 repeats per condition. The stimulus speed and duration were the same as those used in a previous psychophysical study (Kim *et al.*, 2012), which found evidence supporting the claim that humans can integrate audiovisual motion, using a similar type of visual stimulus (random dots) and auditory stimulus (moving bandpass noise). In some penetrations, we also presented an incongruent audiovisual stimulus condition, in which the auditory stimulus moved in the opposite direction to the visual stimulus.

Auditory stimuli were synchronized to the visual stimulus using an AudioFile (Cambridge Research Systems). The AudioFile maintained a set of auditory stimuli as files on internal storage, and was triggered to play a given auditory stimulus when it received a digital signal. We configured the monitor to send a digital signal when the first frame of the visual stimulus appeared, and the AudioFile played the corresponding auditory stimulus within a few hundred microseconds, thereby producing highly synchronized audio-visual stimuli. The selection of the audio stimulus to be played was communicated from the stimulus generation PC to the AudioFile by a USB based DAQ device (USB1208FS, Measurement Computing) before the trial began.

### Determination of receptive fields and basic direction tuning

Visual receptive fields were quantitatively mapped using a grid of either static flashed squares or small apertures of briefly presented moving dots. Visual stimuli were presented full screen, so as to cover as many neurons’ receptive fields as possible. We also conducted visual direction tuning tests, which aided in identifying the location of MT and MST. Direction selectivity was determined using a Rayleigh test (p<0.05) (Berens, 2009).

### Data Analysis

*Time windows and inclusion criteria:* Mean firing rates were calculated using a time window starting 10 ms after stimulus onset (to avoid a potential noise artifact from the speakers) and finishing at the stimulus offset. Units were deemed responsive if they passed a Wilcoxon rank sum test (p<0.01) and if the firing rate was at least 2 spikes/s above the spontaneous rate. Units were considered left-right selective if the firing rate to the best direction of motion (left or right) at 100% coherence was significantly greater than that to the other direction (Wilcoxon rank sum test, p<0.05). Spike rate density functions were calculated using the Chronux software package (Mitra and Bokil, 2008, chronux.org), using a Gaussian kernel with a standard deviation of 25ms. For the population averages analysis, the spike rate density function of each unit was normalized by subtracting the spontaneous firing rate and dividing by the peak spike rate produced during the preferred direction of visual motion at 100% coherence (i.e. the maximum firing rate produced by the unit). These normalized spike rate density functions were then averaged to produce a population normalized spike rated density function for each stimulus type.

*Evoked potentials:* Evoked potentials were calculated by averaging the voltage across trials. We also calculated an average evoked potential for each penetration (across electrodes) to confirm the presence of auditory and visual evoked potentials.

*Neurometric thresholds:* We employed ideal observer analysis to test whether the addition of a moving auditory stimulus improved the ability of neurons to discriminate the direction of motion in a visual left-right discrimination task (Britten *et al.*, 1992). For each level of coherence, we calculated the area under the Receiver Operator Characteristic (aROC) curve from the distributions of responses to the preferred and null directions, the former of which was determined by the direction, either left or right, which elicited the best response at 100% coherence. The aROC values were fitted using least squares regression with two variants of the Weibull function, resulting in a neurometric curve that described the neuron’s performance with respect to coherence (an aROC plot of an example neuron is shown in Figure 5A):

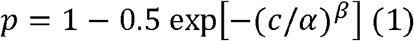

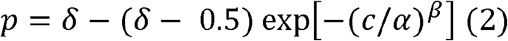

where *p* was the probability of correctly discriminating the direction of motion at coherence *c*, α was the coherence of threshold performance (p=0.82, convention established by Britten et al., 1992), β controlled the slope and δ was the asymptotic level of performance (less than 1). As Equation 2 has an extra free parameter, we used an F-test to decide whether to reject the use of Equation 2 over Equation 1. The α was limited to between 0 and 3, β was limited to lie between 0 and 10, and δ was limited to lie between 0 and 1. Units that did not have an aROC of at least 0.82 at 100% coherence could not have a threshold (i.e. p(c=100)<0.82), were excluded from analyses of thresholds, as was any neuron whose threshold exceeded 100% (given that curving fitting does not guarantee the function will fit all data points perfectly). Statistical testing for differences in the visual and the audio-visual condition was performed with a permutation test (1000 iterations). Each iteration, the visual and audio-visual spike rates were randomly shuffled, and the aROCs and thresholds were recalculated. A distribution of differences in the visual and audio-visual thresholds was constructed from these iterations. A unit was considered to have a statistically significant difference in visual and audio-visual thresholds if the 95% interval of this distribution did not overlap with zero.

*Neurometric performance:* For units that did not meet the above criteria but were still left-right selective, we calculated a “neurometric performance”, which is the integral of the fitted Weibull function from 0 to 100% coherence. This produces a value ranging from 0.5 (uninformative for all coherences) to 1 (perfectly informative across all coherences). Statistical significance testing was performed in the same way as for the neurometric thresholds.

*Population decoding:* For each penetration with at least two left-right selective units, we trained a classifier using Linear Discriminant Analysis to decode the direction of motion at each coherence. We estimated the accuracy and variability of the classifier by training on a randomly sampled subset of 80% (48/60) of trials and testing on the remainder, and repeating this process 1000 times. For each iteration, the population neurometric threshold was calculated in the same way as for the individual units, and the final threshold was calculated as the average threshold across iterations. Differences in visual and audio-visual thresholds were deemed statistically significant if the confidence interval of the distribution of threshold differences did not overlap zero.

### Histology

At the end of the recordings, the animals were given an intravenous overdose of sodium pentobarbitone (100 mg/kg) and, following cardiac arrest, were perfused with 0.9% saline, followed by 4% paraformaldehyde in 0.1 M phosphate buffer pH, 7.4. The brain was post-fixed for approximately 24 hours in the same solution, and then cryoprotected with fixative solutions containing 10%, 20%, and 30% sucrose. The brains were then frozen and sectioned into 40 μm coronal slices. Alternate series were stained for Nissl substance and myelin (Gallyas, 1979). The location of recording sites was reconstructed by identifying electrode tracks and depth readings recorded during the experiment. Additionally, each electrode array was coated in DiI, allowing visualization under fluorescence microscopy prior to staining of the sections (Figure 1C). In coronal sections, MT is clearly identifiably by heavy myelination in the granular and infragranular layers (Figure 1C) (Rosa & Elston, 1998), whereas MST is more lightly myelinated and lacks clear separation between layers (Palmer & Rosa, 2006a). The majority of units reported here were histologically confirmed to be in MT or MST, but for some penetrations in which the histology was unclear (12% of units), units were included on the basis of their receptive field size and progression, and their direction tuning.

## Results

### Sample size

We made 27 electrode array penetrations in areas MT and MST (MT: n=18; MST: n=7; MT/MST: 2), and recorded 314 visually responsive units (MT: n=223; MST: n=91), of which 11% were classified as single-units. We did not find any significant difference between single and multi-units in the following analyses, so they were grouped together. In agreement with previous reports (Zeki, 1974; Maunsell & Van Essen, 1983; Albright, 1984; Celebrini & Newsome, 1994; Born & Bradley, 2005; Lui & Rosa, 2015), we found that most units showed visual direction selectivity (MT 76% MST 60%, Rayleigh test, p<0.05). To test for audio-visual integration, we only presented leftwards or rightwards motion, but did not restrict our analyses to units which preferred one of these directions. We found that 238 (76%) of the units we recorded showed significantly different responses to visual leftwards versus rightwards motion (Wilcoxon rank sum test, p<0.05; MT: n=180; MST: n=58).

### Auditory stimuli do not elicit spiking responses in MT/MST

In all penetrations, we presented moving visual stimuli as well as moving and stationary auditory stimuli. Although we recorded a large number of visually responsive units, we did not observe any spiking responses to any of the auditory stimuli. Figure 2 shows the spiking activity of 4 example units that were selective for visual motion on the left-right axis, and one unit which was not left-right selective. Visual and audiovisual stimuli evoked strong spiking responses over time for at least one direction of motion at 100% coherence (blue and red lines, bottom and middle rows of the rasters). In contrast, moving and stationary auditory stimuli didnot evoke responses at all (green lines, top row of rasters).

**Figure 2:**
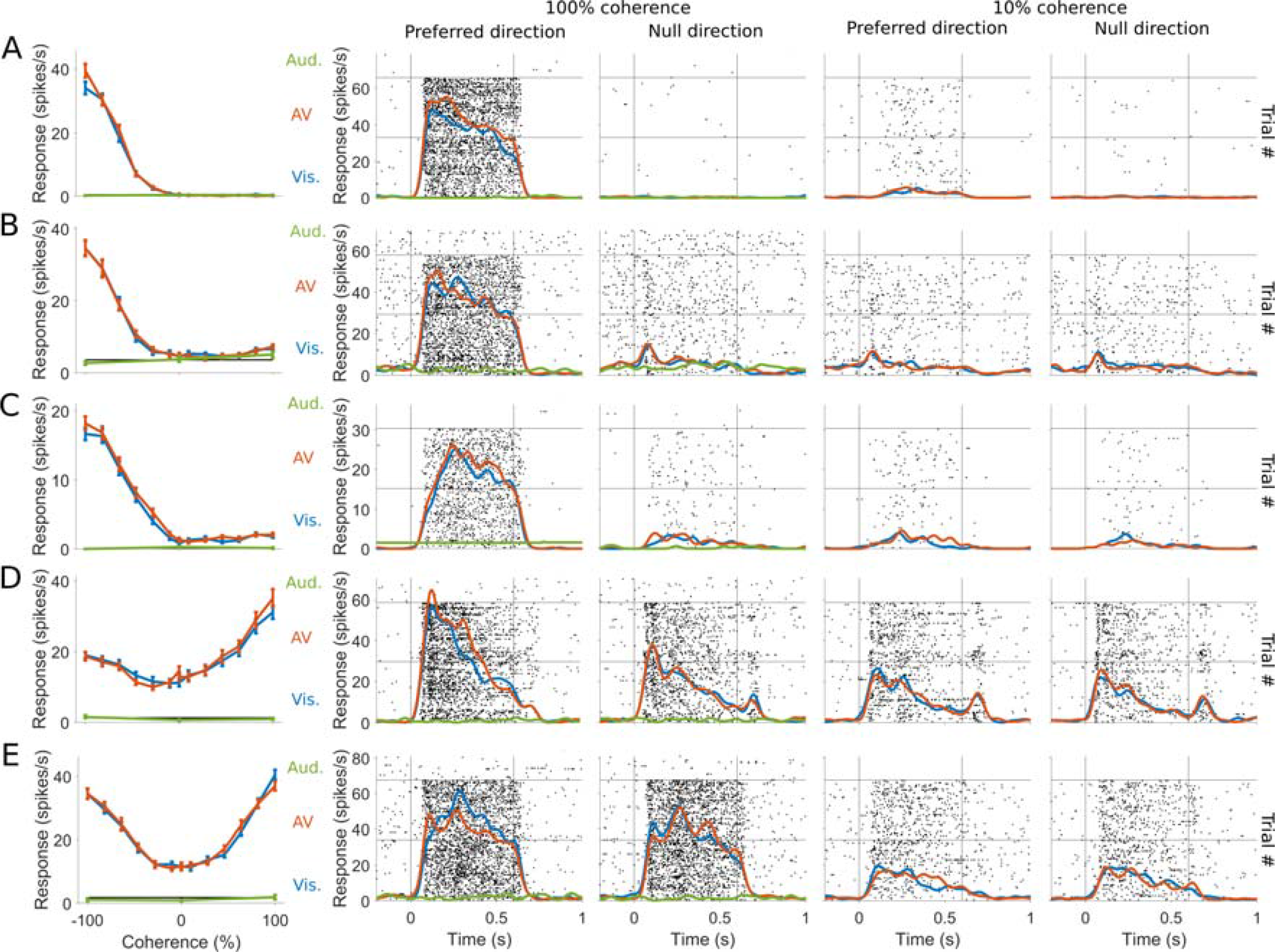
Example units showing clear and robust responses to visual but not auditory stimuli. Each row (A-E) shows a different unit - A: single-unit in MST, B: multi-unit in MST, C: single-unit in MT, D & E: multiunits in MT. In each row, the first panel shows the mean firing rate in response to leftwards or rightwards motion. For visual and audiovisual motion, the level of dot coherence was manipulated, as shown by the x-axis, with leftwards motion represented by negative values, and rightwards motion represented by positive values. For the auditory stimuli, only 3 conditions were used - leftwards, rightwards and stationary (plotted as responses to -100, 100 and 0% coherence respectively). The remaining panels show the spiking responses to motion in the preferred and null directions at 100% and 10% coherence, with the spike rate density functions overlaid on top of raster plots. The raster plots are separated into visual, audiovisual and auditory conditions by the horizontal lines, as shown the by the legend in the first column (note there were fewer trials for auditory only stimuli, and no equivalent of 10% coherence for auditory stimuli). The vertical lines show the onset and offset of the stimuli.

Figure 3 shows the distributions of the mean firing rates of the best moving visual and auditory stimuli (panels A and B, respectively) in 3 animals in which we interleaved visual and auditory motion trials (n=203 units). None of the visually responsive units showed statistically significant responses to auditory stimuli (determined by a Wilcoxon Rank Sum test, p<0.01). To confirm that the auditory stimulus which we used could evoke spiking activity elsewhere in the brain, we recorded from auditory areas adjacent to MST as a positive control (Figure 1A, dark green region) and found units that produced clear responses to the moving auditory stimulus (e.g. Figure 3C). There were also small but discernable evoked potentials in response to auditory click stimuli, even in MT and MST (Figure 3D), confirming that auditory stimulation evoked neuronal activity in all animals. However, the MT/MST auditory evoked potential was small in comparison to the visual (Figure 3D, red vs blue trace). The auditory cortex is within 4 mm of our recording sites in areas MT and MST in the marmoset (Figure 1A), so it is likely that this auditory evoked potential was the result of activity in distal auditory brain regions (most likely caudal auditory areas: Palmer and Rosa, 2006a; Kajikawa and Schroeder, 2011) rather than being generated by neurons in MT/MST.

**Figure 3:**
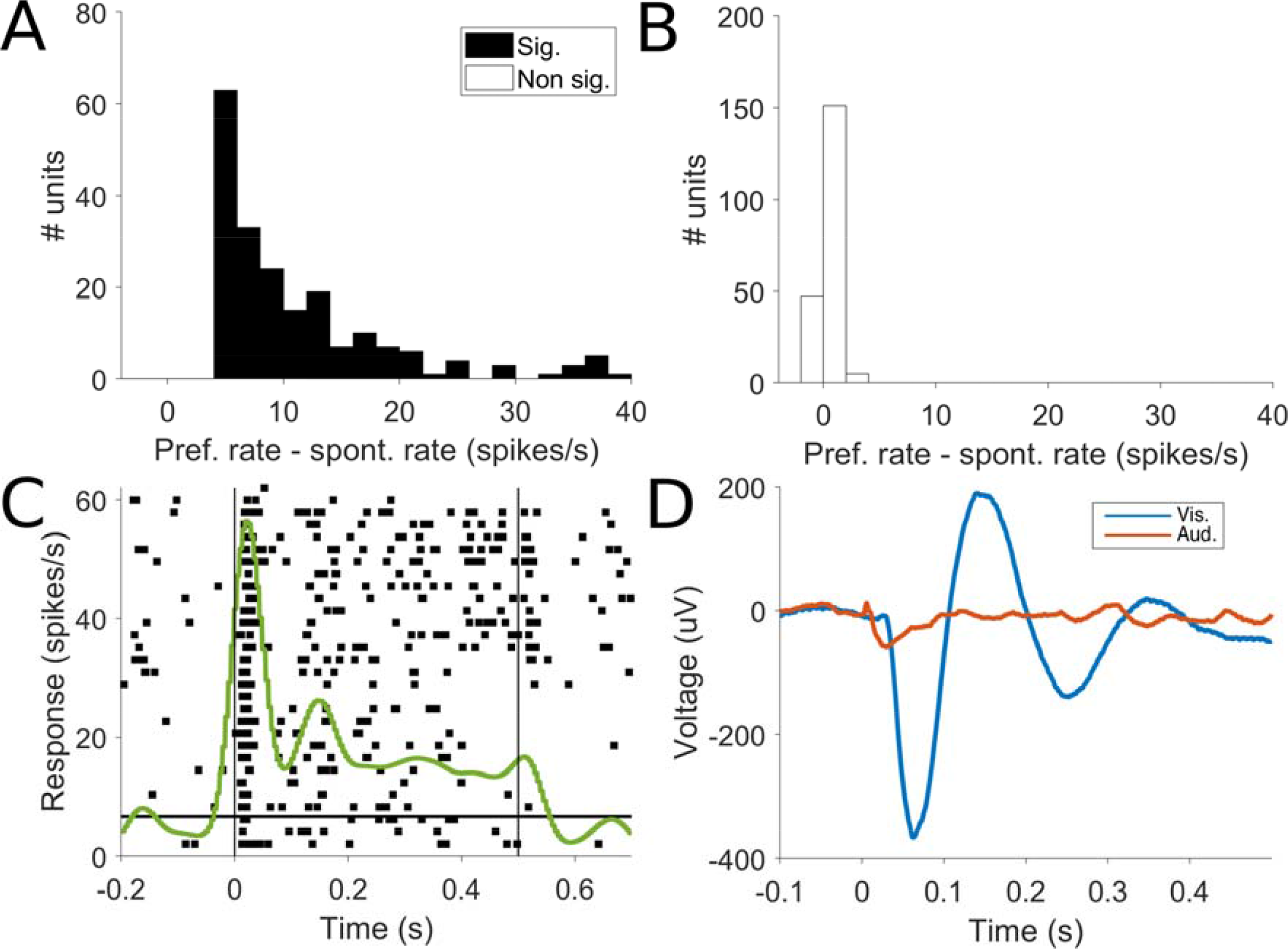
Responses to visual and auditory motion. A: Firing rates of visually responsive units to the best direction of motion (left or right) from 3 animals in which visual and auditory stimuli were interleaved in the same testing block. By definition, all units show statistically significant different evoked rates compared to the spontaneous rate, and are therefore colored black, as indicated in the legend. B: Responses of the same units in A to the best direction of auditory motion (left or right). None of these responses were significantly different from the spontaneous rates, and are colored white to denote this. C: An example raster plot and spike density function of multi-unit activity in the auditory cortex in response to auditory motion, demonstrating that this stimulus can evoke spiking activity. D: Visual flash and auditory click evoked potentials, averaged across all visually responsive channels from all animals.

### Moving auditory stimuli do not modulate firing rates in MT/MST

As moving and static auditory stimuli did not elicit spiking activity, we investigated if the addition of a moving auditory stimulus modulated the firing rate of neurons to a moving visual stimulus in the same direction. We tested this at different levels of motion coherence, given that multisensory integration is more likely to occur when the unimodal signal strength (or neural response) is weak (Meredith & Stein, 1983; Meredith *et al.*, 1987; Deneve *et al.*, 2001; Ma *et al.*, 2006; Gu *et al.*, 2008; Fetsch *et al.*, 2011). Neurons in areas MT and MST are known to be direction selective (Dubner & Zeki, 1971; Felleman & Kaas, 1984; Celebrini & Newsome, 1994), but as we only presented motion along the horizontal axis, we observed a range of responses to changes in coherence (Figure 2). Some units showed a response pattern consistent with a clear preference for one direction (e.g. Figure 2A-C) (Britten *et al.*, 1992), some had only a moderate degree of left-right selectivity (e.g. Figure 2D), and others showed no preference for leftwards or rightwards motion (e.g. Figure 2E). The majority of the last grouping (68%) were direction selective for visual motion but preferred directions of motion close to the vertical axis.

To analyze the full population, each unit was designated as right-preferring or left-preferring based on which direction elicited the highest mean spike rate at 100% coherence, with the preferred direction being designated as positive 100% coherence, and the null direction as negative 100% coherence. To visualize the population response, the responses of each unit were first normalized to lie between 0 and 1, so that 0 is the spontaneous rate and 1 is the response to the preferred direction, and then averaged to produce a population-average response to motion in the preferred and null directions, at different levels of coherence, for visual-only and audio-visual conditions (Figure 4A). A 2-way repeated measures ANOVA did not find a significant effect of modality (visual or AV; F_1,11_=4.41×10^−4^, p=0.983), only coherence (F_1,11_=88.2, p=8.06×l0^−189^), on firing rates, and did not find a significant interaction effect between modality and coherence (F_1,11_=0.007, p=1). We also found that there was no difference in firing rates due to modality when units from MT and MST were analyzed separately (2-way ANOVA, F_1,11_=4.73×10^−6^, p=0.998, and F_1,11_=0.0031, p=0.956, respectively for main effect of modality). To test if auditory stimuli had any effect on the 0% coherence stimulus, we used a 1-way ANOVA to test if there was any difference in firing rate in the visual-only condition compared to the audiovisual conditions with auditory motion in the preferred or null visual direction, but again found no effect (F_2_=0.416, p=0.660).

**Figure 4.**
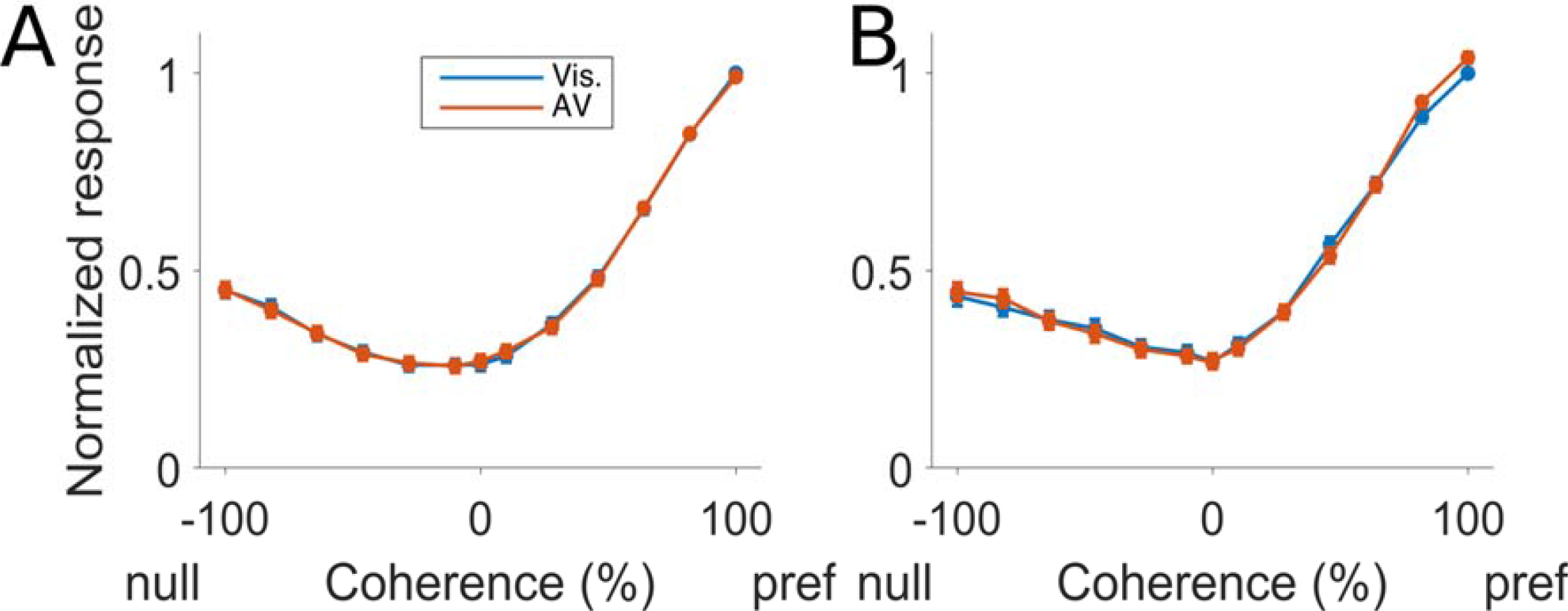
A: Normalized population coherence response functions for visual and audio-visual response. Error bars show the standard error of the mean. Motion in the preferred direction is represented by positive coherence values and motion in the null direction is represented by negative coherence values. B: Same convention as A, showing a subset of units in which the auditory component of the audiovisual motion moved in the opposite direction to the visual stimulus (i.e. the two modalities were incongruent).

MT receptive fields can have a complex centre-surround organization that enhances motion contrasts (Allman *et al.*, 1985). We tested the possibility of an antagonistic interaction between visual and auditory motion by using incongruent audiovisual motion, in which the auditory stimulus moved in the opposite direction to the visual motion (Figure 4B). In a subset of units in which we performed this test (n=203), we did not observe any difference in firing rates between the visual and the incongruent audiovisual condition (2-way repeated measures ANOVA, modality F_1,11_=0.014, p=0.906, coherence F_1,11_=128, p=1.62×10^−259^, interaction F_1,11_=0.223, p=0.996), or when MT and MST where analyzed separately (MT n=162: modality F_1,11_=0.0304, p=0.862, coherence F_1,11_=83.6, p=5.31×10^−170^, interaction F_1,11_=0.258, p=0.992; MST n=41: modality F_1,11_=0.0125, p=0.911, coherence F_1,11_=47.7, p=2.75×10^−83^, interaction F_1,11_=0.145, p=1). In summary, there was no evidence to suggest that auditory motion modulates the spike rates of MT/MST neurons, and this was consistent for all levels of visual motion noise tested.

### Moving auditory stimuli do not improve visual neurometric thresholds

The analysis in the previous section included all visually responsive units, even those that do not carry information regarding the direction of motion (e.g. Figure 2E). Working on the hypothesis that auditory stimuli may enhance motion discriminability or sensitivity, we analyzed just the responses of units that show strong left-right selectivity (e.g. Figure 2A-C). We used aROC analysis to determine how well an ideal observer could determine the direction of motion (left or right) from the firing rate of a neuron (Newsome *et al.*, 1989; Britten *et al.*, 1992) at different levels of motion coherence, to determine whether the addition of the auditory stimulus resulted in improved neurometric performance compared to the visual only stimulus. We calculated the aROC at each coherence and fitted a Weibull function (e.g. Figure 5A) to determine the neurometric threshold - the coherence level at which the aROC reaches 0.82 (by convention, Britten et al., 1992, see inset of Figure 5A). Of the 238 left-right direction selective units recorded, 135 (57%; MT: n=103; MST: n=32) reached a maximum aROC of at least 0.82, and thus had a defined neurometric threshold. The mean change in neurometric threshold for the audio-visual versus visual conditions was not statistically significant different from zero (Figure 5B, 0.3% coherence impairment for the audio-visual condition, paired t-test p=0.88). We did not find any difference when units from MT and MST were analyzed separately (p=0.83 and p=0.94 respectively, paired t-tests). Only 3 units (2.2%, permutation test p<0.05) showed statically significant differences (Figure 5B, black bars); we regard these as likely false positives.

**Figure 5:**
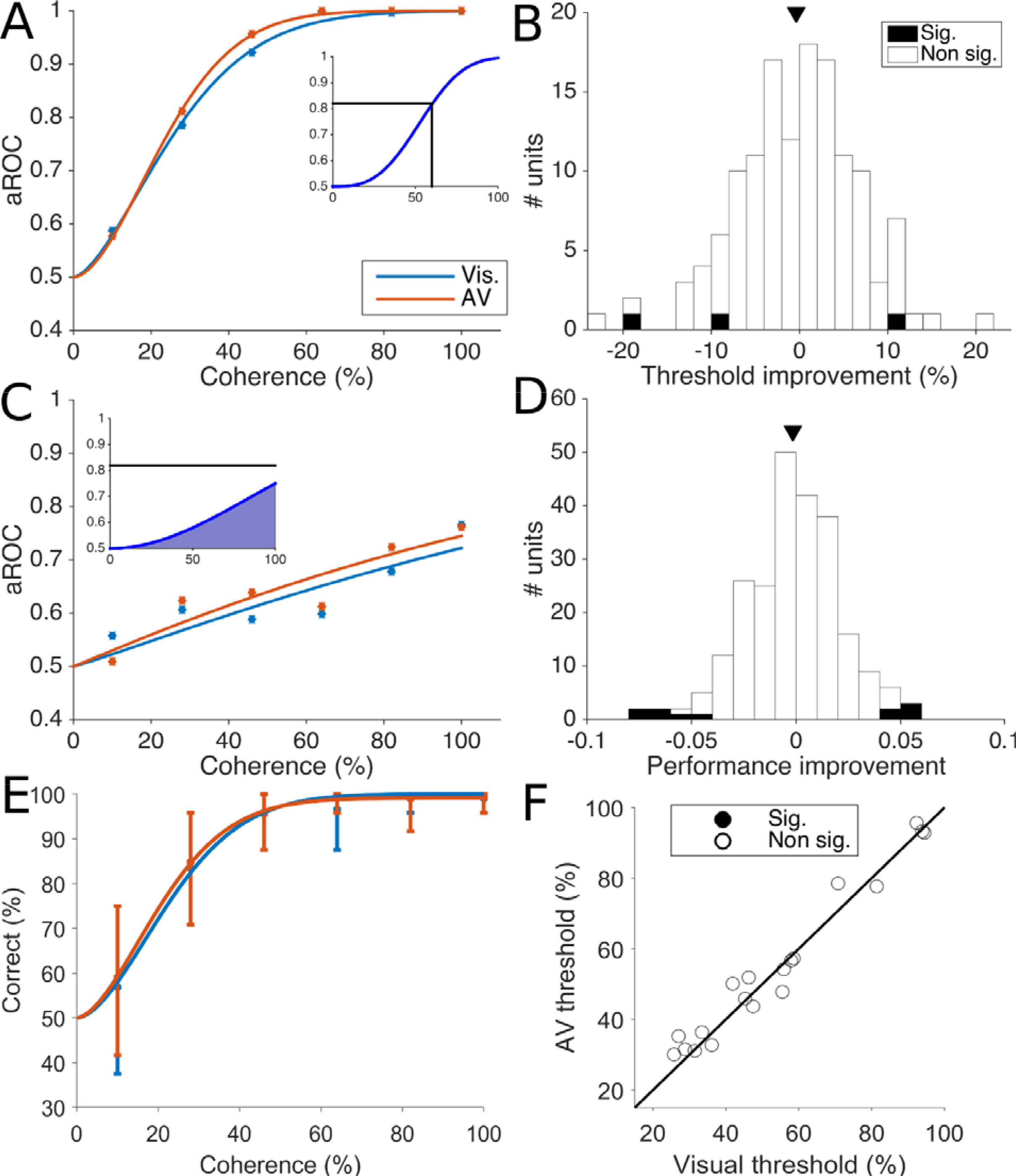
Neurometric thresholds and performance. A: Example neurometric curves for the unit from Figure 2A. Visual neurometric curve and the audio-visual neurometric curve are shown. The inset shows how the neurometric threshold is calculated (the coherence level that results in an aROC of 0.82). B: Distribution of the improvements of the audio-visual versus visual neurometric threshold, with the mean shown by the triangular marker at the top. Significance for individual units is shown as per the legend. C: Example neurometric functions for the unit from Figure 2D, which does not have a defined neurometric threshold (i.e. does not reach an aROC of 0.82). The inset shows how the neurometric performance was calculated (the area under the curve). D: Distribution of the improvement of the audio-visual versus visual neurometric performances; the mean is shown by the triangular marker, and individual units showing significant differences are indicated by bar color. E: Example population decoding from one penetration. This penetration contained 25 individual units. The x-axis shows the coherence level, and the y-axis shows the performance of the classifier, with error bars representing the 95% interval of cross-validation iterations. F: Comparison of visual (x-axis) and audio-visual (y-axis) population neurometric thresholds. Individual penetrations showing significant differences are colored as indicated by the legend, none were significantly different.

Of the left-right selective units, 103 (43%) did not meet the criteria for threshold analyses. To investigate the contribution of all left-right selective units, we measured the neurometric performance as the area under the fitted Weibull curve, as this does not require that a unit achieves any particular threshold aROC. Larger areas correspond to better discrimination performance across coherences (e.g. Figure 5A vs Figure 5C). In line with the neurometric threshold analyses, we found that the mean change in neurometric performance for the audiovisual versus visual condition was not significantly different from zero (Figure 5D, 0.002 unit impairment for the audio-visual condition, paired t-test p=0.86). Again, there was no difference when units from MT and MST were analyzed separately (p=0.75 and p=0.9 respectively, paired t-tests). Only 11 (4.6%) of the units showed a statistically significant difference between the visual and audio-visual conditions (permutation test, Figure 5D, black bars), and the effects were balanced in terms of whether the audio-visual stimulation improved or decreased performance.

Trial to trial correlations between neurons can have profound effects on population coding (Bair *et al.*, 2001; Cohen & Newsome, 2009; Quian Quiroga & Panzeri, 2009; Ruff & Cohen, 2016; Panzeri *et al.*, 2017) and so we tested if auditory stimulation affects the encoding of stimuli at the population level. To investigate this, we trained a classifier to decode the direction of motion (left or right) for each of the 25 array penetrations that contained at least 2 left-right selective units. Of these, 21 penetrations had defined population neurometric thresholds (analogous to the individual unit neurometric thresholds - performance of at least 82% at 100% coherence). Figure 5E shows the performance of one such penetration, containing 25 units from area MT. We measured the neurometric performance of each penetration under visual and audio-visual conditions (Figure 5F), but found none that showed significant differnces in visual and audio-visual thresholds. Moreover, the mean difference in visual and audio-visual thresholds across penetrations was not significantly different from zero (Figure 5F; 0.11% coherence improvement in the audio-visual condition, p=0.98, paired t-test). In summary, the addition of the moving auditory stimli in a congruent direction to random dot visual stimuli did not increase or decrease the amount of information carried by single neurons or populations of simultaneously recorded neurons.

### Temporal response profile for visual, auditory and audiovisual stimuli

To investigate whether there were any modulations of spiking activity in specific time ranges that may not have been detected in the previous analysis (which counted spikes over the full stimulus duration), we examined the temporal profile of spiking activity in response to visual, auditory and audio-visual stimuli. To test whether the addition of auditory stimulus enhanced (or diminshed) directional information contained in spiking neurons, we analysised the the subset of units (n=135) that had defined neurometric thresholds and were therefore strongly left-right selective (Figure 5A,B), we calculated normalized spike rate density functions for each direction and level of motion coherence (Figure 6). Confirming the findings from Figure 3, the mean firing rate in response to auditory stimuli was not significantly different to to zero for all time bins during the stimulus presentation (25ms time bins, t-test, p < 0.01), demonstrating there was no brief or transient response to the auditory stimulus at any stage of the stimulus presentation. In agreement with the analysis in Figures 4 and 5, the mean firing rate of the visual and audiovisual conditions was not significantly different at any time bin during the stimulus presentation (25ms time bins, t-test, p < 0.01). Therefore auditory responses and audio-visual modulations were not present at any time point in the population of the strongly left-right selective units.

**Figure 6:**
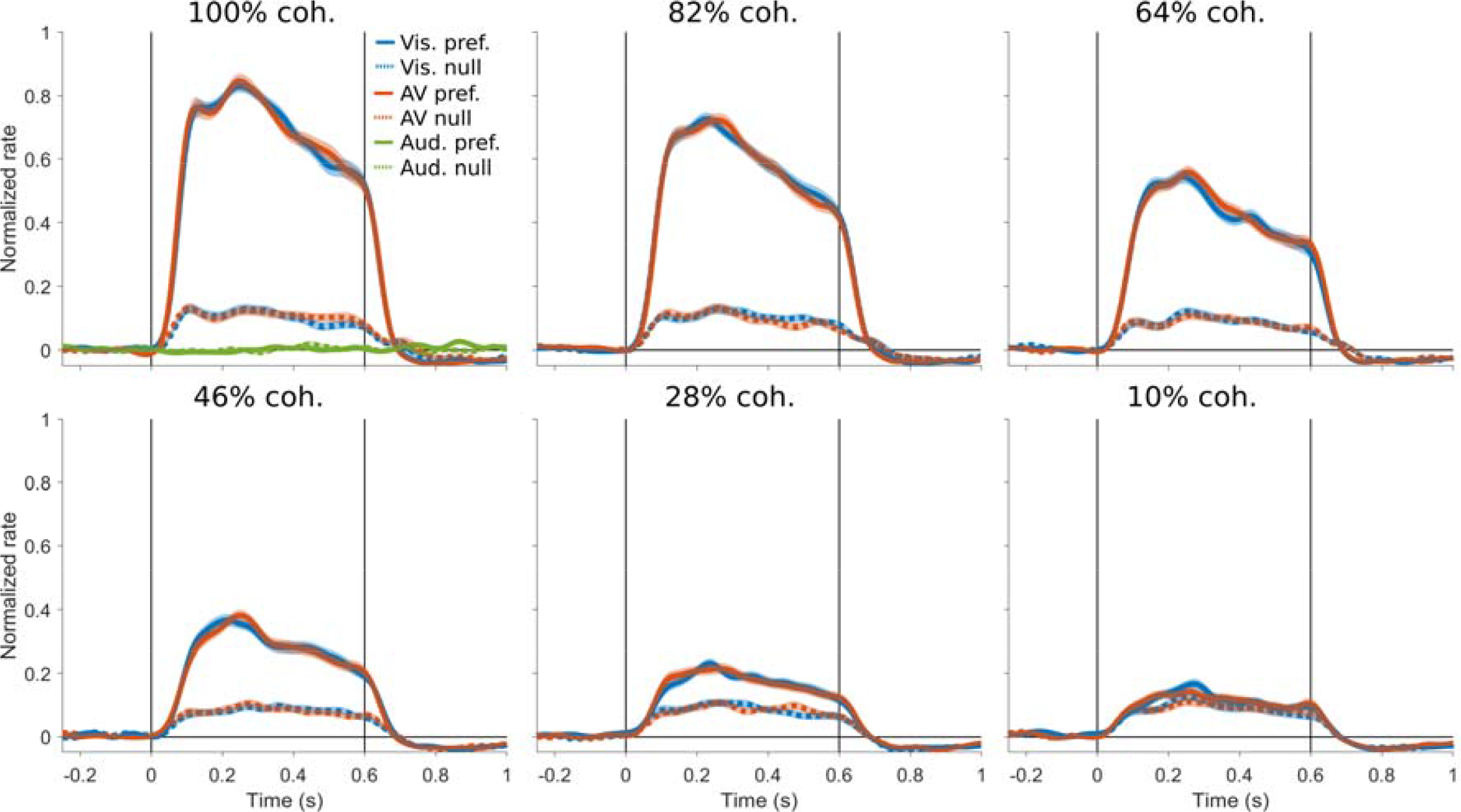
Temporal response profile of spiking activity in response to visual, auditory and audio-visual stimuli. The normalized population-averaged spike rate density function for each direction and coherence is shown in a separate panel, with the standard error of the mean shown by the shading. The response to visual, audio-visual and auditory stimuli in the preferred and null direction of motion are colored as indicated by the legend in the first panel. Only the 100% coherence panel shows auditory responses since there was no equivalent of coherence (stimulus noise) for the auditory stimuli. The spiking response is not modulated by the auditory stimuli at any time and the difference between the responses to the visual and audiovisual stimuli was not different at any time point.

## Discussion

We investigated the extent to which neurons in the visual motion processing areas MT and MST respond to auditory motion stimuli, and whether congruent auditory stimuli modulate the activity of these neurons in conjunction with visual motion. Using an anaesthetized preparation in which both auditory and visual responses can be robustly elicited, we found no evidence of cross-modal interactions in spiking activity in either MT or MST neurons. Neither stationary nor moving auditory stimuli elicited spiking when presented alone, and moving auditory stimuli did not modulate firing rates when paired with visual stimuli. We investigated this using methods for characterizing the information carried by both individual neurons (Britten *et al.*, 1992; Gu *et al.*, 2008) and for populations of neurons. We found no evidence of multi-sensory integration at either near-threshold or supra-threshold levels of stimulus noise (Meredith & Stein, 1983; Deneve *et al.*, 2001; Ma *et al.*, 2006; Gu *et al.*, 2008; Fetsch *et al.*, 2011).

### Experimental conditions affecting the lack of audio-visual integration

As our study was conducted using an anesthetized preparation, it could be argued that we did not observe cross-modal effects because they are only present in the conscious state, or when the animal is performing an audio-visual motion discrimination task. Although we cannot rule out this explanation, it should be noted that multisensory responses have been widely reported under anesthesia in areas of the cortex corresponding to multiple hierarchical levels (Bruce *et al.*, 1981; Hikosaka *et al.*, 1988; Wallace *et al.*, 1992; Bizley *et al.*, 2007), as well as in subcortical structures (Meredith & Stein, 1983; Reig & Silberberg, 2014). Therefore multisensory integration can be supported, at least in part, by bottom-up processes which are still present in the anesthetized state (Alkire *et al.*, 2008); in fact, multisensory responses in the superior colliculus can be hard to detect in the awake state because of motor related activity (Bell *et al.*, 2001, 2003). Our results indicate that the integration of audio-visual motion cues is not likely to be supported by bottom-up integration in areas MT and MST. However, the possibility remains that multisensory responses may be evident in the awake state if they are supported by inputs from high-order polysensory association areas, such as the superior temporal polysensory cortex (area TPO; Baylis *et al.*, 1987; Boussaoud *et al.*, 1990), and if these inputs are selectively affected by anaesthesia. In the superior colliculus, multisensory integration (but not multisensory responses) is dependent on descending inputs from the cortex (Alvarado *et al.*, 2009), even under anesthesia.

It is also possible that audio-visual integration would have been evident if different types of stimuli were used. However, MT neurons respond well, and in a direction-selective manner, to many types of visual motion (bars: Maunsell & Van Essen, 1983; Albright, 1984; Lui *et al.*, 2012; gratings: Movshon *et al.*, 1985; dot patterns: Britten *et al.*, 1992; Solomon *et al.*, 2015; Chaplin *et al.*, 2017) and a relatively broad range of speeds. We thus regard as unlikely that MT cells would require very specific auditory stimuli to activate, or integrate, which were not covered within the frequency and ILD range of our broadband stimuli.

Our experiments required the use of headphones to simulate auditory motion by manipulating the ILD, but it could be argued that demonstrating multisensory responses in MT and MST necessitates other auditory motion cues, such as interaural time differences and pinnae-based spectral cues. However, similar set-ups using headphones have been used in human studies that have shown audio-visual integration for motion (Kayser *et al.*, 2017), and ILDs have been shown to be the dominant cue for sound localization in the azimuth for marmosets for high frequency sounds (Slee & Young, 2010). Yet, we observed no auditory responses (Figure 2) or modulations (Figures 3 and 4) when manipulating the ILD. In summary, the auditory stimuli used in the present study contained strong cues for sound location and motion, which would be expected to elicit at least observable changes in neural activity should MT and MST neurons receive this information.

### Comparison to human studies

Using single and multiunit recordings, we did not observe responses to moving auditory stimuli in either MT or MST neurons, a result that reflects the conclusions of several imaging studies in humans (Bedny *et al.*, 2010; Alink, Euler, Kriegeskorte, *et al.*, 2012; Jiang *et al.*, 2014). Two other studies have reported activation in hMT+ in response to moving auditory stimuli (Poirier et al., 2005; Strnad et al., 2013), but it has been suggested that this may be explained by the methodology that was used to localize hMT+, and that the auditory-related effects may arise from a cortical area outside hMT+ (Jiang *et al.*, 2014). Our observations are compatible with the latter point of view.

Other fMRI studies have shown that moving auditory stimuli can modulate hMT+ activity when paired with visual stimuli (Lewis & Noppeney, 2010; Strnad *et al.*, 2013; von Saldern & Noppeney, 2013). One key difference between these studies and ours is that the human participants are conscious and often performing a task. Therefore the audio-visual related activity may not correspond to audio-visual cue integration itself, but to task-related activity (Alink, Euler, Kriegeskorte, *et al.*, 2012; Bizley *et al.*, 2016; Kayser *et al.*, 2017), such as the binding of the two modalities to form a unified percept (Nahorna *et al.*, 2012, 2015; Bizley & Cohen, 2013), attentional effects (Bavelier *et al.*, 2000; Beer & Roder, 2004; Beer & Röder, 2005; Lakatos *et al.*, 2008), or choice-related signals from the decision making process. Indeed, it is well known that top-down attention can affect spiking activity in MT and MST (Treue & Maunsell, 1996) and this extends to feature based attention, which is selective for direction of motion (Treue & Martinez-Trujillo, 1999).

Another key difference is that the hemodynamic response observed in fMRI, as well as the evoked potentials observed in EEG, are likely to reflect synaptic (input) activity rather than spiking (output) activity (Logothetis *et al.*, 2001; Buzsáki *et al.*, 2012). We also observed auditory evoked potentials in MT and MST to auditory click stimuli (Figure 3D). Intracranial auditory evoked potentials have been reported in the visual cortex of mice (Iurilli *et al.*, 2012; Olcese *et al.*, 2013; Ibrahim *et al.*, 2016), and recently in humans (Brang *et al.*, 2015). Thus, it may be possible that areas MT and MST do receive auditory information, as suggested by studies which demonstrate that they receive sparse inputs from the auditory cortex (Palmer & Rosa, 2006a, 2006b). However, we could not rule out the possibility that these evoked potentials we observed in MT and MST originated from nearby auditory cortex, which is only a few millimeters away from our recording sites (Palmer & Rosa, 2006a; Kajikawa & Schroeder, 2011). As we did not find any evidence of auditory modulation of visually evoked spiking in response to motion, it is unlikely that any auditory inputs to MT or MST lead directly to audio-visual motion cue integration.

### Implications for multisensory integration in the cerebral cortex

Overall, our results favor the traditional model of cortical multisensory processing, with separate cortical domains for each modality, and multisensory neurons being restricted to intermediate zones between these domains (e.g. Avillac *et al.*, 2007; Foxworthy *et al.*, 2013). Most of the physiological evidence for auditory influences on spiking activity in visual cortex have come from studies in mice (Iurilli *et al.*, 2012; Olcese *et al.*, 2013; Meijer *et al.*, 2017), which have direction connections between the primary visual and primary auditory cortex (Meredith & Lomber, 2017), reflecting a general scaling rule whereby smaller brains tend to show more direct connectivity across cortical systems (Horvát *et al.*, 2016; Rosa *et al.*, 2018). Direct anatomical connections between the visual and auditory cortex in primates have been reported (Falchier *et al.*, 2002; Rockland & Ojima, 2003; Cappe & Barone, 2005; Palmer & Rosa, 2006a, 2006b), but to our knowledge, there is only one study that showed that auditory stimuli can modulate the spiking activity in the visual cortex of primates (Wang *et al.*, 2008), producing a modest decrease in response latency in awake animals. On the other hand, cross modal influences in auditory cortex have been observed across species (primates: Lakatos *et al.*, 2007; ferrets: Bizley *et al.*, 2007; Bizley & King, 2009; Meredith & Allman, 2015).

Whether or not MT/MST participates in multisensory integration is also likely to be modality dependent: it is well established that area MST (but not MT: Chowdhury *et al.*, 2009) responds to and integrates vestibular motion cues, with studies showing clear evidence of spiking responses (Duffy, 1998; Gu *et al.*, 2006) and multisensory integration (Gu *et al.*, 2008; Fetsch *et al.*, 2011). Other human imaging studies have indicated that hMT+ is involved in processing tactile stimuli (Hagen *et al.*, 2002; Beauchamp *et al.*, 2007; Basso *et al.*, 2012; Pei & Bensmaia, 2014), but much like auditory processing (Jiang *et al.*, 2014), these findings remain controversial (Jiang *et al.*, 2015).

## Abbreviations

aROC: area under the receiver operating characteristic curve
AV: audio-visual
BOLD: blood oxygenation level dependent
hMT+: human middle temporal area complex
MST: medial superior temporal area
MT: middle temporal area

## Acknowledgements

We thank Nicolas Price and Rowan Tweedale advice on the manuscript, and Katrina Worthy for assistance with the histological work. We also thank Janssen-Cilag for the donation of sufentanil citrate. This project was funded by the National Health and Medical Council of Australia (LL: APP1066232) and the Australian Research Council (LL: DE130100493; MR: CE140100007). TC was funded by an Australian Postgraduate Award.

## Conflict of interest

The authors declare no conflict of interest.

## Author contributions

TC, LL, MR and RR designed the research; TC, BA, MH, RR and LL performed the research; TC analyzed the data; TC, LL, RR and MR wrote the paper

## Data statement

Data will be made available on request to the corresponding authors.

